# Diffusion in a quantized vector space generates non-idealized protein structures and predicts conformational distributions

**DOI:** 10.1101/2023.11.18.567666

**Authors:** Yufeng Liu, Linghui Chen, Haiyan Liu

## Abstract

The power of diffusion probabilistic models (DDPMs) in protein design was recently demonstrated by methods that performs three-dimensional protein backbone denoising. However, these DDPMs tend to generate protein backbones of idealized secondary structures and short loops, lacking diverse, non-idealized local structural elements which are essential for the rich conformational dynamics of natural proteins. Moreover, the sampling power of DDPMs have not yet been utilized for predicting the conformational distributions of natural proteins of dynamic structures. Aiming at these two needs, we developed a model named PVQD (protein vector quantization and diffusion), which used an auto-encoder with vector quantization and a generative diffusion model in the latent space to jointly performing the challenging task of modeling complicated protein structures within an end-to-end framework. Our study demonstrated that in design PVQD generated designable protein structures containing non-idealized elements, while in single sequence-based structure prediction PVQD reproduced experimentally observed conformational variations for a set of natural proteins of dynamic structures.

## Introduction

In past years, deep learning methods have made significant progresses in protein structure prediction ^1, 2^ and protein design ^3, 4, 5, 6^. Translational and rotational invariant or equivariant deep networks have been invented for joint and end-to-end modeling of sequential data and three-dimensional data, yielding unprecedented accuracy in structure prediction based on multiple sequence alignments ^1, 2^. Such networks have also been applied in amino acid sequence design for given protein backbones (*i.e*., inverse protein folding) ^3, 4, 7^ and *de novo* protein backbone design (*i.e*., protein structure generation) ^8, 9, 10, 6^. Several resulting models have been demonstrated to achieve superior experimental success rates and design capabilities over conventional methods ^3, 4, 6^.

Despite these developments, some important needs in protein structure prediction and protein design are still unsatisfied. One is the reliable sampling of structure distributions due to conformational dynamics. This need has not been fulfilled with existing end-to-end methods for protein structure prediction that use deterministic networks ^11, 12, 13, 14^. Another unfulfilled need is the design of protein structures that possess preferred conformational dynamics. Protein structures designed with current methods are biased to contain idealized secondary structure elements (mostly helices) and short loops ^9, 10, 6^. Such structures, although designable and stable, are likely to be highly rigid, lacking conformational dynamics essential for functions such as such as catalysis or allosteric regulations.

In principle, generative networks based on sampling unconditional or variously conditioned distributions including the denoising diffusion probabilistic models (DDPMs) ^15, 16, 17^ can be adopted to unify the tasks of conformation sampling and *de novo* structure design. DDPMs of protein structures reported so far either performed diffusion directly in the structural space ^8, 9, 10, 6^, or used non-end-to-end approaches in which structural restraints developed from diffusion models were applied to conventional energy function-based modeling ^18^. A limitation of diffusion directly in the structural space is that the complicated and strong correlations between the high-dimensional structural variables need to be captured by the diffusion model alone, which could be too demanding for current DDPMs based on noise of Gaussian distributions. While the non-end-to-end approaches can be less demanding with the diffusion models, these approaches inherit the limitations in accuracy and computational efficiency of energy function-based modeling. With these limitations, the ability to design experimentally realizable protein structures by most reported DDPMs have not yet been confirmed ^8, 10, 18^, while the few exceptional DDPMs supported by experimentally determined protein structures still tend to generate structures of idealized local elements and high rigidity ^9, 6^. Moreover, a DDPM that can be conditioned on single amino acid sequence to sample the conformational distributions of natural proteins is yet to be described ^19^.

In this work, we developed a model named PVQD (protein vector quantization and diffusion), in which diffusion was not performed in the original structure space but in a latent representation space of residue-wise three-dimensional structural contexts. This representation was learnt through auto-encoding with vector quantization. Compared with DDPMs in the three-dimensional structure space, the main advantage of PVQD is to divide the challenging task of end-to-end modeling of complicated protein structures between an auto-encoder and a diffusion model. We demonstrated that PVQD unified structure design and prediction in a single framework. We demonstrated that in design PVQD generated designable protein structures composed of non-idealized structure elements, while in prediction PVQD reproduced experimentally observed conformational variations for a set of natural protein.

## Results

### An overview of PVQD

The overall architecture of PVQD is depicted in Fig. 1. Details of the various network blocks are presented in Methods. Briefly, the method comprises an auto-encoder with vector quantization and a generator using denoising diffusion.

**Fig 1.**
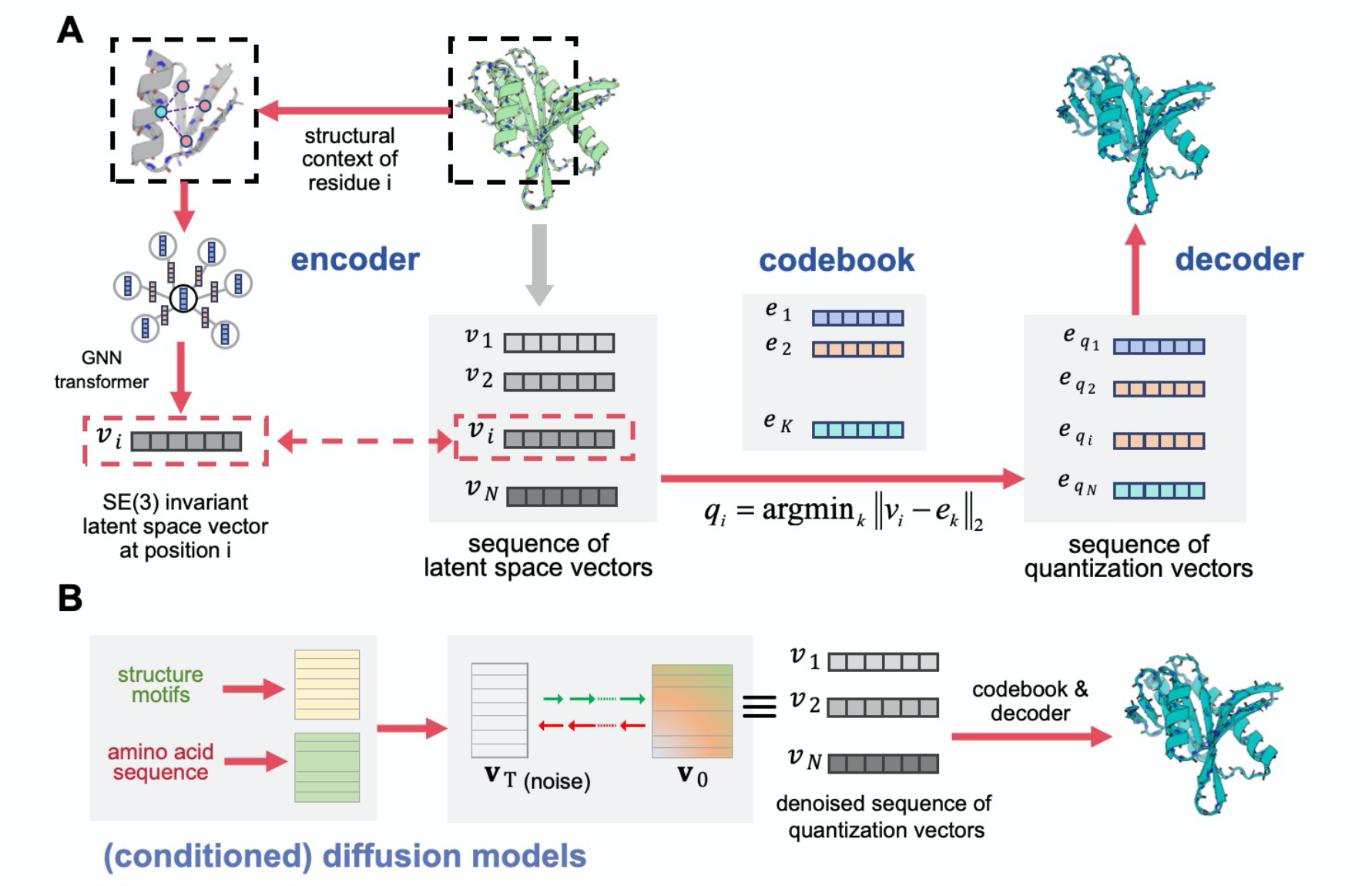
The overall architecture of PVQD. A. The auto-encoder of PVQD comprises an encoder, a codebook and a decoder. The encoder transforms the three-dimensional backbone structure into a sequence of SE(3)-invariant latent space vectors. These vectors are replaced by the closest quantization vectors contained in the codebook. The resulting sequence of quantization vectors are used by the decoder to best recover the original three-dimensional backbone structure. The generator of PVQD performs denoising diffusion in the latent vector space to generate denoised sequences of quantization vectors, from which three-dimensional backbones structures are constructed through the codebook and decoder. The generator can be controlled by conditioners encoding structural restraints (for structure design) or amino acid sequences (for structure prediction).

The auto-encoder (Fig. 1A) learns to represent the structural context of an amino acid residue in a protein backbone structure as a discrete code. It consists of an encoder, a codebook, and a decoder. Using features describing the relative geometries between a central residue and its spatially neighboring residues as input, the encoder generates a translational and rotational invariant (*i.e*., SE(3)-invariant) vector in a latent space to embed this residue-wise structural contexts. The codebook maps the latent vector space to a finite set of discrete codes (the structure codes) by using vector quantization. The decoder reconstructs the overall three-dimensional structure from the sequence of the quantization vectors (deduced from the sequence of structure codes or the code sequence) produced by the encoder and the codebook. Together with the structure decoder, an auxiliary residue type decoder (not displayed in Fig. 1A) recovering residue types at different sequence positions from the sequence of the quantization vectors is employed to enable the use of amino acid sequence recovery losses as extra regularization terms during model training.

For structure generation, PVQD uses diffusion based on noise of Gaussian distributions to model and sample from the joint distribution of the latent space vectors of all residues (Fig. 1B). The sampled sequence of quantization vectors is mapped to a sequence of the quantization vectors and decoded into three-dimensional structures with the pre-trained codebook and decoder from the auto-encoder. Following ideas used for the conditional generation of images ^20^, conditional structure generation with PVQD is developed by coupling embedded conditions with latent space diffusion. The conditions of given structural restraints and of given amino acid sequence are implemented with respective conditioner modules, which are coupled with downstream diffusion modules. The different conditioned diffusion models are obtained from the unconditioned one through fine-tuning.

### The accuracy of the auto-encoder of PVQD

We evaluated the accuracy of the PVQD auto-encoder on a set of non-redundant natural protein structures selected with the PISCES server ^21^ (sequence identity cutoff 40% and X-ray structure resolution cutoff 2.0 Å). We measured the structural deviations of the reconstructed backbones from corresponding input backbones with the template modeling scores (recTM-scores) and root mean square deviations (recRMSDs) of atomic positions, with higher recTM-scores and smaller recRMSDs indicate high reconstruction accuracy. Figures 2A and 2B show the distributions of these metrics for proteins divided into different groups according to their sizes. All the recTM-scores are above 0.42; for proteins of less than 450 residues, the recRMSDs are distributed below 1.5 Å with the median values of all groups below 1.0 Å; and for proteins of 450 to 850 residues, the median recRMSD values are below 3.0 Å. Fig. 2C shows an example of a protein of more than 600 residues, displaying a superimposition of the reconstructed backbone with the original natural structure. It demonstrated that both the secondary structure elements and the loops were reconstructed with high accuracy.

**Fig 2.**
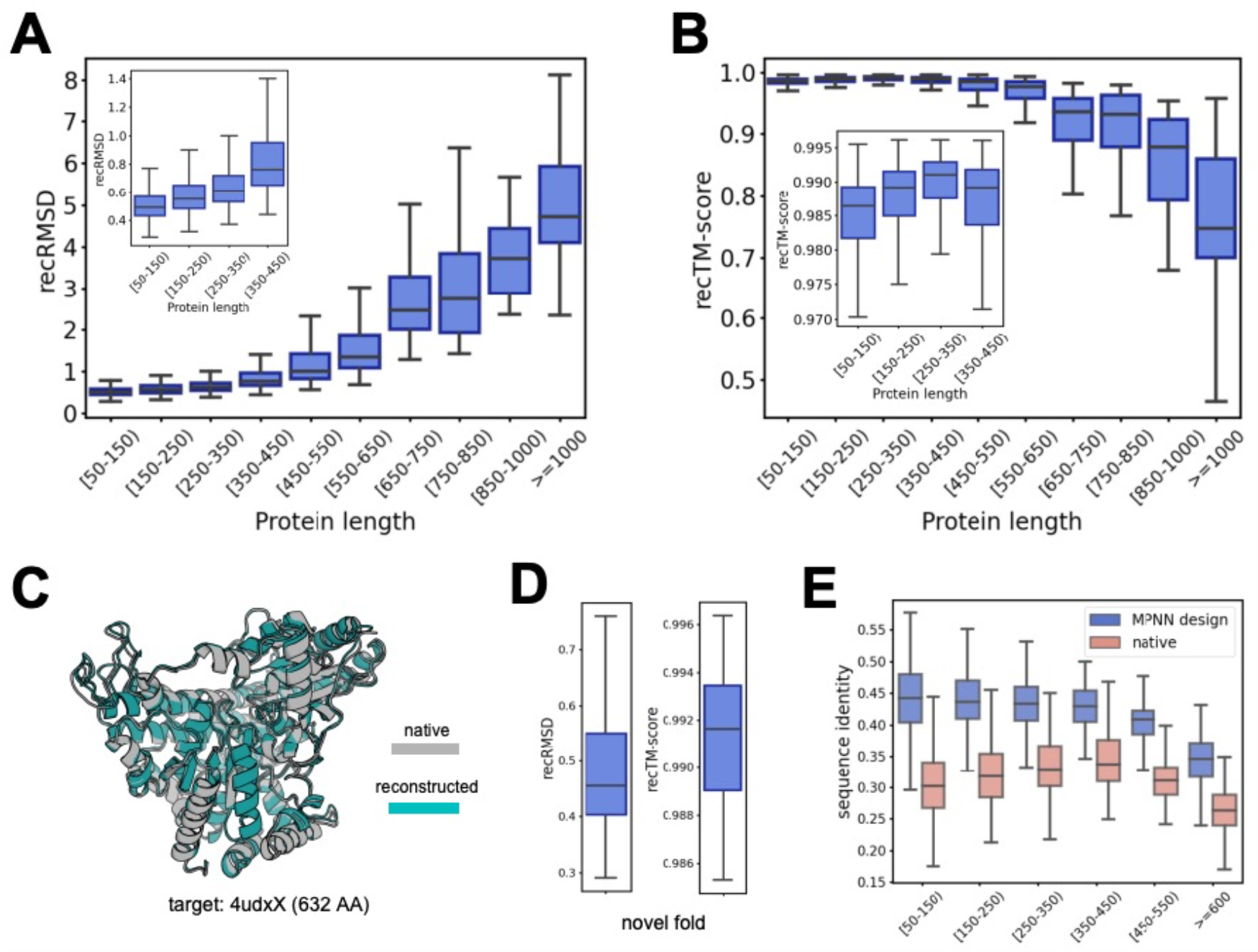
The accuracy of the auto-encoder of PVQD. A. Distributions of the RMSDs (recRMSD) between the backbones reconstructed with the PVQD auto-encoder and the corresponding natural backbones. B. Distributions of the TM-scores (recTM-scores) between the backbones reconstructed with the PVQD auto-encoder and the corresponding natural backbones. C. A reconstructed protein backbone (green) with more than 600 residues superimposed with the corresponding natural backbone (gray). The PDB ID of the natural backbone is indicated. D. Distributions of the recRMSDs and recTM-scores for the set of non-natural backbones generated by SCUBA-D. These backbones are of dissimilar overall folds to natural backbones with highest TM-scores to PDB below 0.5 and designable with self-consistent RMSDs (scRMSDs) below 2.0 Å. E. Distributions of the identities of the amino acid sequences decoded by the auxiliary amino acid type decoder with the native amino acid sequences (blue) and with the ProteinMPNN-designed amino acid sequences (salmon). The boxplots show median, interquartile range, and minimum and maximum values excluding outliers (>1.5 times the interquartile range beyond the box).

To examine if the auto-encoder can be generalized to structures not seen during training (*e.g*., structures absent from PDB), we considered 63 non-natural, designable backbones generated using SCUBA-D (a diffusion model for generating designable protein backbones) ^9^. These structures are of highest TM-scores to PDB blow 0.5 (which indicate that these structures are of little similarity to PDB structures in overall folds) and self-consistent RMSDs (scRMSDs) below 2.0 Å (which indicate that these structures are designable) (here, the scRMSD refer to the RMSD between a designed backbones and the structure predicted by AlphaFold2 (AF2) ^2^ on designed amino acid sequences based on that backbone. Unless otherwise specified, fixed-backbone amino sequence design was performed with the program ProteinMPNN ^4^). Fig. 2D shows that the auto-encoder reconstructed these backbones accurately with recTM-scores above 0.96 and recRMSDs below 1.5 Å.

For the natural proteins, we examined the identities between the amino acid residue types recovered by the residue type decoder in the PVQD auto-encoder with corresponding native residue types. For most proteins, the identities are above 30% (median value 32%) (Fig. 2E). Interestingly, the recovered residue types are of higher identities (median value 43%) with those designed with ProteinMPNN (a program for designing amino acid sequence based on given backbone) than with the native sequences (Fig. 2E). For the SCUBA-D generated backbones, we used AF2 to predict structures from amino acid sequences composed of the decoded residue types and determined the corresponding scRMSDs. The results showed that only 8 out of 63 scRMSD values were below 2.0 Å. Thus, compared with fixed-backbone design programs such as ProteinMPNN, the auxiliary residue type decoder of PVQD is not a robust amino acid sequence designer, perhaps because it chooses the amino acid types at different positions independently.

### PVQD generates diverse and designable backbones

We generated and evaluated backbones of 70 to 400 residues. The distribution of the mutual TM-scores between the generated backbones are shown in Fig. 3A. The small median TM-score (0.29) suggests that these backbones are diverse, spread across a wide range of structures of dissimilar overall folds. The distributions of the highest TM-scores to PDB shown in Fig. 3B indicate that a substantial number of generated backbones are of low similarity to PDB structures (*e.g*., 23 out of 100 backbones of 200 residues are of highest TM-scores to PDB below 0.5). This suggests that the diffusion generator of PVQD generalizes to protein structures not seen during training.

**Fig 3.**
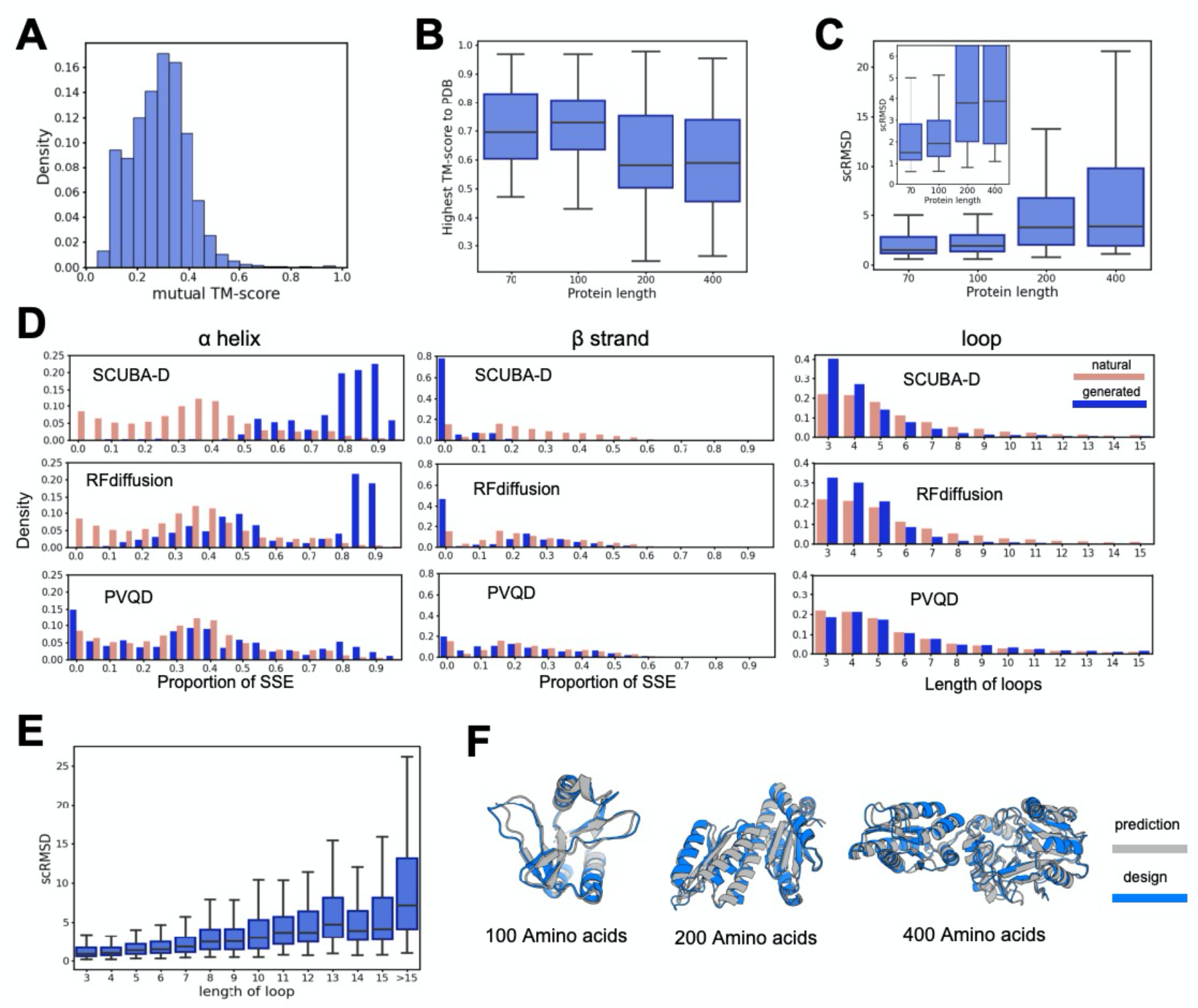
Computational analyses of protein structures unconditionally generated with PVQD. A. The distribution of the mutual TM-scores between the generated backbones of 100 to 400 residues. B. Distributions of the highest TM-scores of the generated backbones to existing structures in the PDB. C. Distributions of the self-consistent RMSDs (scRMSDs), which are the RMSDs between the generated backbones and the structures predicted by ESMFold for amino acid sequences designed with the generated backbones. The analyzed backbones were of 100 to 400 residues and structure predictions were performed on 8 ProteinMPNN-designed amino acid sequences for each backbone. D. The histograms of the proportions of residues in the α helix state (left) and in the β strand state (middle) and the histograms of loop lengths (right) for the unconditionally generated backbones with SCUBA-D (top), RFdiffusion(middle) and PVQD (bottom). For comparisons, the corresponding histograms computed on the structures of a set of non-redundant proteins are also shown. E. The distribution of the scRMSDs of loops between the unconditionally generated backbones and the corresponding predicted backbones. Each scRMSD was calculated by superimposing 4 residues flanking the loop on each side. F. Three examples of the unconditionally generated backbone (blue) superimposed with the corresponding predicted backbone (gray). The boxplots show median, interquartile range, and minimum and maximum values excluding outliers (>1.5 times the interquartile range beyond the box).

It has been noted that diffusion from noise in the structural space is more likely to lead to structures of α-helices than of β-sheets, probably because those helices composed of contiguous sequence segments are easier to form than β-sheets of disparate segments. Compared with structures generated with SCUBA-D ^9^ and RFdiffusion ^6^, the unconditionally generated structures of PVQD showed secondary structure compositions closer to natural proteins in PDB (Fig. 3D). Especially, the probabilities of structures containing high ratios of β-sheets were significantly increased. The distribution of loop lengths in PVQD generated structures are also closer to that observed in natural proteins. Some structures contain long loops of 15 residues or more while such loops are rarely found in *de novo* protein structures designed with previous methods. coordinates diffusion model.

Visual inspections of the backbones generated by PVQD suggest that these backbones are often composed of multiple structural domains of sizes distributed similarly as in natural proteins. This is different from RFdiffusion or SCUBA-D, which tend to generate large single domain structures even for chain lengths of up to 400 to 500 residues (Fig. 4). This may have caused PVQD to be more likely to generate backbones that fold similarly as natural proteins (according to the distributions of the highest TM-scores to PDB, Fig. 3C) than RFdiffusion or SCUBA-D for the unconditional generation of backbones of more than 200 residues. We note that the multi-domain backbones are also likely to be of larger scRMSDs than the single domain backbones, as the inter-domain arrangements of the designed backbone and the backbone predicted from the designed amino acid sequence may differ even when the individual domains were of small scRMSDs.

**Fig 4.**
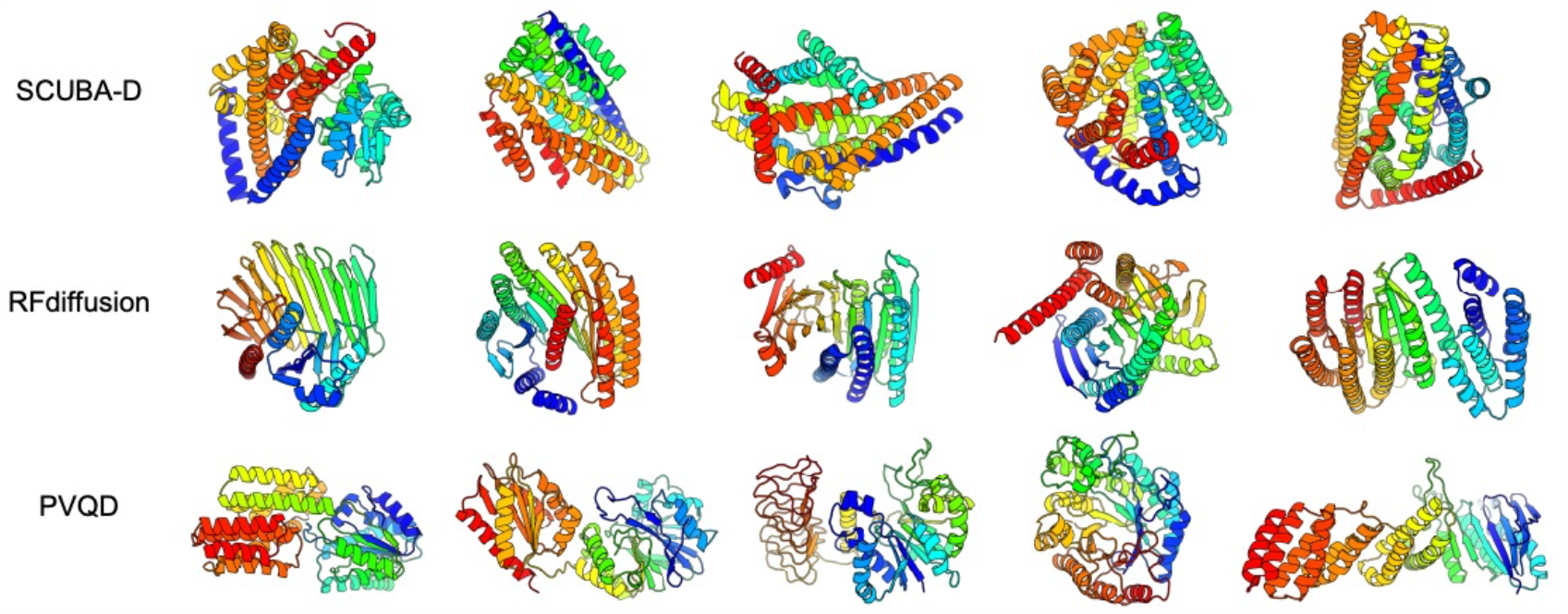
Examples of 400-residue backbones unconditionally generated with different methods. Top: SCUBA-D; middle: RFdiffusion; and bottom: PVQD.

### Designing backbones satisfying structural restraints with PVQD

*De novo* backbone generators like PVQD can be applied to design functional proteins through motif scaffolding, in which certain parts of a generated backbone are restrained to form a predefined functional motif (*i.e*., a pre-defined partial structure comprising amino acid residues responsible for an intended function). We implemented such restraints with conditional diffusion in PVQD. We evaluated the resulting model by using a benchmark set containing 17 motif-scaffolding problems from simple structure “inpainting” to the scaffolding of motifs including viral epitopes, receptor traps, ligand interfaces, small molecule binding sites, and enzyme active sites. Previously, this benchmark set was used to evaluate the Evodiff ^22^ and the RFdiffusion methods. As in that study, we generated 100 backbones for each problem, applied ESMFold ^23^ to predict the structures for amino acid sequences designed on the generated backbones. and counted a problem as being successfully “solved” when at least one predicted structure for the problem has a motifRMSD (*i.e*., the RMSD of atomic positions between the motif residues in the predicted structure and those in the pre-defined motif structure)below 1.0 Å and a pLDDT score above 70.

PVQD found acceptable solutions for 14 problems, slightly higher than the results of Evodiff (12 problems) and RFdiffusion (13 problems) (see Fig. 5A). Interestingly, PVQD and RFdiffusion found solutions for the same 13 problems (see Fig. 5B). The median value of the success rates for individual problems of RFdiffusion was higher than that of PVQD, especially for those problems in scaffolding the motifs consisted of less than 5 residues. For these problems, the RMSDs between the PVQD generated motif backbones and the desired motif backbones were already large (see Fig, 5C). This suggested that the current PVQD generator was not yet able to use a partial structural context defined by only a few residues as proper conditions.

**Fig 5.**
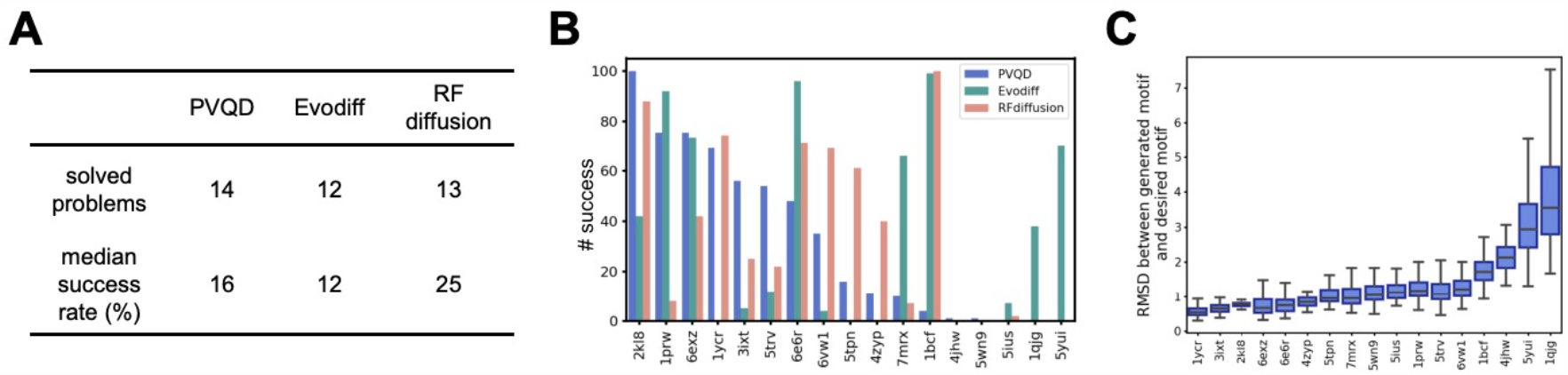
The performance of PVQD in solving the motif-scaffolding problems. A. The numbers of problems solved by different methods. We counted a problem as being successfully “solved” when at least one predicted structure for the problem has a motifRMSD below 1.0 Å and a pLDDT score above 70. The total number of problems in the benchmark set is 17. B. The success rates for each problem of the three methods: PVQD (blue), Evodiff (green) and RFdiffusion (salmon). C. The distributions of the backbone RMSDs between the motif structures in the PVQD-generated backbones and the desired motif structures.

### Predicting conformational distributions of natural proteins with PVQD

We preformed single amino acid sequence-based structure prediction on several commonly considered benchmark sets with PVQD and compared with previously reported results for other models (Table 1). PVQD exhibited at least comparable, and mostly better performances than other single sequence-based methods for these benchmark sets ^19, 24^.

**Table 1.**
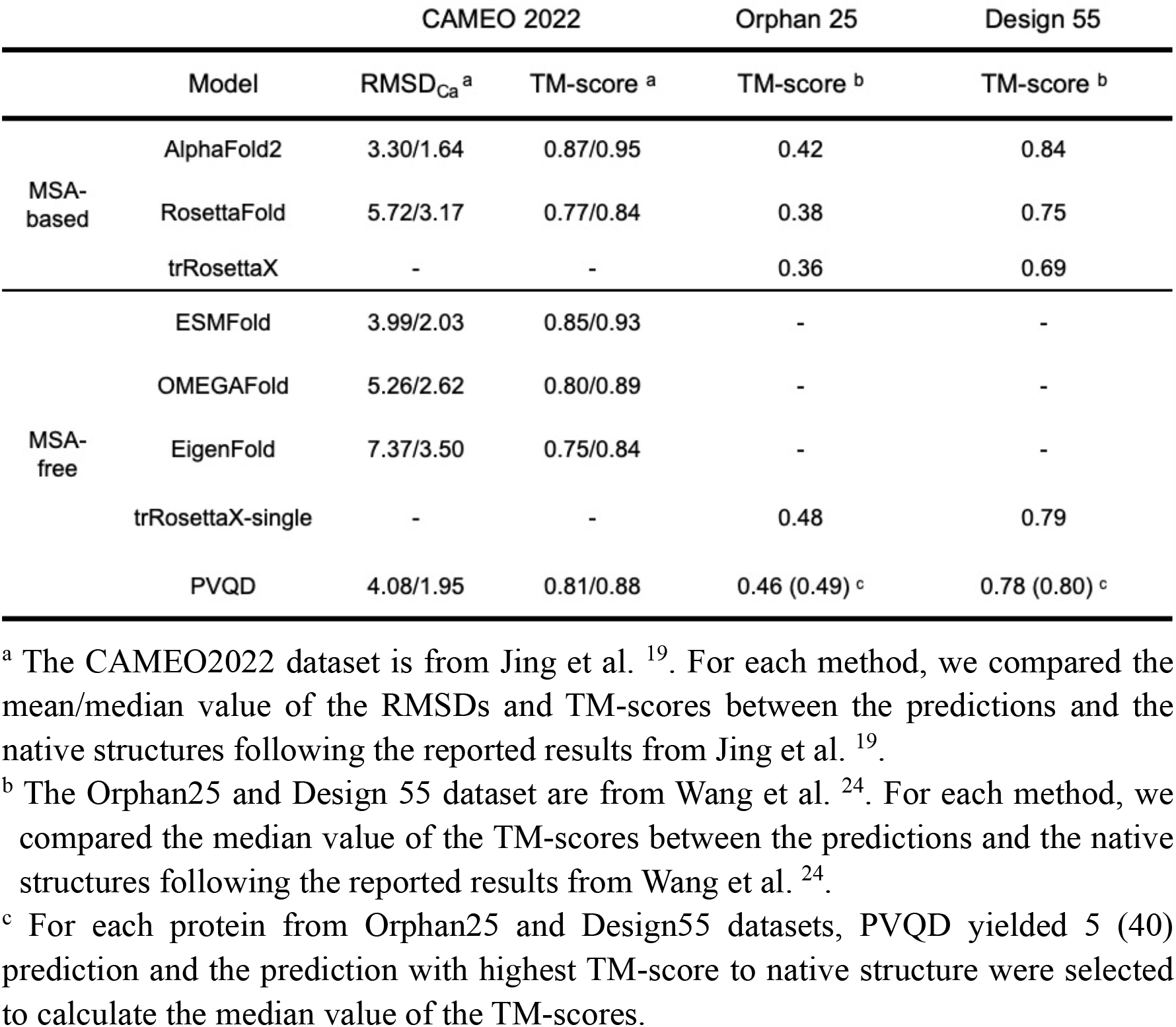
The performances of PVQD in structure prediction compared with other methods for three benchmark sets.

|  |  | CAMEO 2022 |  | Orphan 25 | Design 55 |
| --- | --- | --- | --- | --- | --- |
|  | Model | RMSD<sub>Ca</sub><sup>a</sup> | TM-score<sup>a</sup> | TM-score<sup>b</sup> | TM-score<sup>b</sup> |
| MSA-based | AlphaFold2 | 3.30/1.64 | 0.87/0.95 | 0.42 | 0.84 |
|  | RosettaFold | 5.72/3.17 | 0.77/0.84 | 0.38 | 0.75 |
|  | trRosettaX | - | - | 0.36 | 0.69 |
| MSA-free | ESMFold | 3.99/2.03 | 0.85/0.93 | - | - |
|  | OMEGAFold | 5.26/2.62 | 0.80/0.89 | - | - |
|  | EigenFold | 7.37/3.50 | 0.75/0.84 | - | - |
|  | trRosettaX-single | - | - | 0.48 | 0.79 |
|  | PVQD | 4.08/1.95 | 0.81/0.88 | 0.46 (0.49)<sup>c</sup> | 0.78 (0.80)<sup>c</sup> |

<sup>a</sup> The CAMEO2022 dataset is from Jing et al. <sup>19</sup>. For each method, we compared the mean/median value of the RMSDs and TM-scores between the predictions and the native structures following the reported results from Jing et al. <sup>19</sup>.
<sup>b</sup> The Orphan25 and Design 55 dataset are from Wang et al. <sup>24</sup>. For each method, we compared the median value of the TM-scores between the predictions and the native structures following the reported results from Wang et al. <sup>24</sup>.
<sup>c</sup> For each protein from Orphan25 and Design55 datasets, PVQD yielded 5 (40) prediction and the prediction with highest TM-score to native structure were selected to calculate the median value of the TM-scores.

Next, we evaluated the conformational ensembles sampled with PVQD by considering a benchmark set of 91 natural proteins for each of which at least two experimentally determined alternative structures were available ^19^. For each protein, we predicted 40 backbone structures with PVQD, and identified for each experimental structure of the protein the predicted structure of the highest TM-score. For 74 proteins, the resulting highest TM-scores were above the TM-scores between the corresponding alternative experimental structures (Fig. 6A). Fig. 6B displayed the corresponding TM-score distributions by considering proteins of X-ray structures and for proteins of NMR separately. Fig. 6C displayed the TM-score distributions for proteins exhibiting different types of structural changes (based on manual inspection): localized conformational changes (the “local” group); inter-domain motions (the “inter-domain” group); and structural changes that are not typically local changes or inter-domain motions (the “other” group). The highest prediction-versus-experiment TM-scores distributed in higher ranges than the TM-scores between experimental structures. This suggests that the errors of approximating different experimental structures with different predicted structures are on average smaller than the amplitudes of the experimentally observed structural changes.

**Fig 6.**
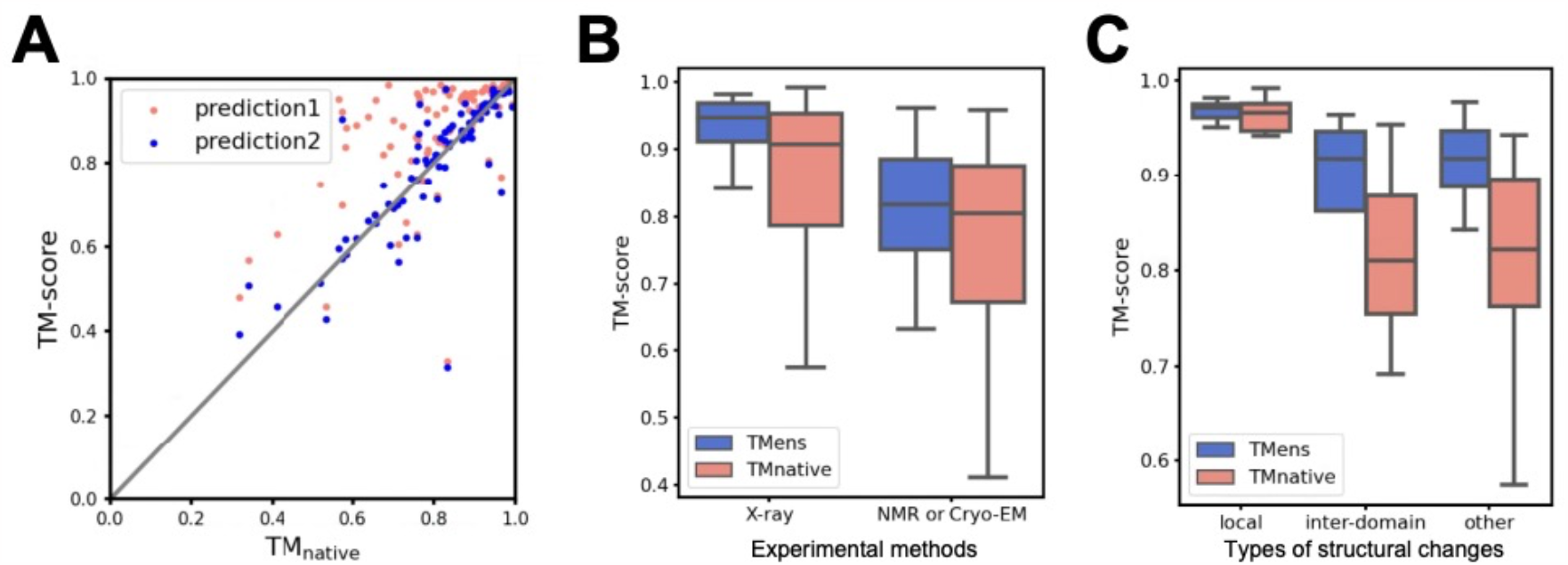
Comparisons of the PVQD-predicted structures with the experimental structures for a set of proteins with multiple conformations. A. For each protein, two experimental structures and 40-PVQD predicted structures were considered. For each experimental structure, the predicted structure of the highest TM-score was identified. The corresponding highest TM-scores (*i.e*., ensemble TM-scores) versus the TM-scores between the experimental structures of the same proteins (*i.e*., experimental structure TM-scores) were shown in the scattering plot. For each protein, the point representing higher prediction-versus-experiment TM-score is colored in salmon while the point representing lower prediction-versus-experiment TM-score is colored in blue. B. Distributions of the mean ensemble TM-scores (colored in blue) and of the experimental structure TM-scores (colored in salmon). The proteins were grouped according to the experimental methods used for determining their structures. C. The same as B, but the proteins were grouped according to their types of structural changes.

We further considered three example proteins displaying different types of structural changes in their experimental structures. For each protein, we predicted 2000 conformations with PVQD, visualized the predicted structures on a two-dimensional plane (obtained with Multi-Dimensional Scaling or MDS ^25^) together with the experimental structures and AF2-predicted structures (with pLDDT scores above 80). The PVQD-generated structures were clustered with the K-means algorithm and a representative structure for each cluster was found ^26^.

The predicted structure for K-Ras, a protein displaying localized structural changes between different experimental structures ^27, 28^, formed three main clusters on the 2-D MDS plane (see Fig. 7A). In Fig. 7B, superimpositions of the representative structure of cluster 1 with the representative structures of clusters 2 and 3 are displayed. All 20 experimental structures of K-Ras fall into cluster 1 on the 2-D MDS plane. The RMSDs of the experimental structures from the respectively most similar predicted structures (the smallest RMSDs) were from 0.68 to 1.41 Å, with the smallest RMSDs for 14 experimental structures being below 1.0 Å (see Fig. 7C). Fig. 7D show the superimpositions of the experimental and predicted structure pair with the lowest smallest RMSD and of the pair with the highest smallest RMSD. Fig. 7E shows that there are high correlations between the residue-wise root-mean-square fluctuations (RMSFs) of the predicted structures in cluster 1 and the same RMSFs between the experimental structures.

**Fig 7.**
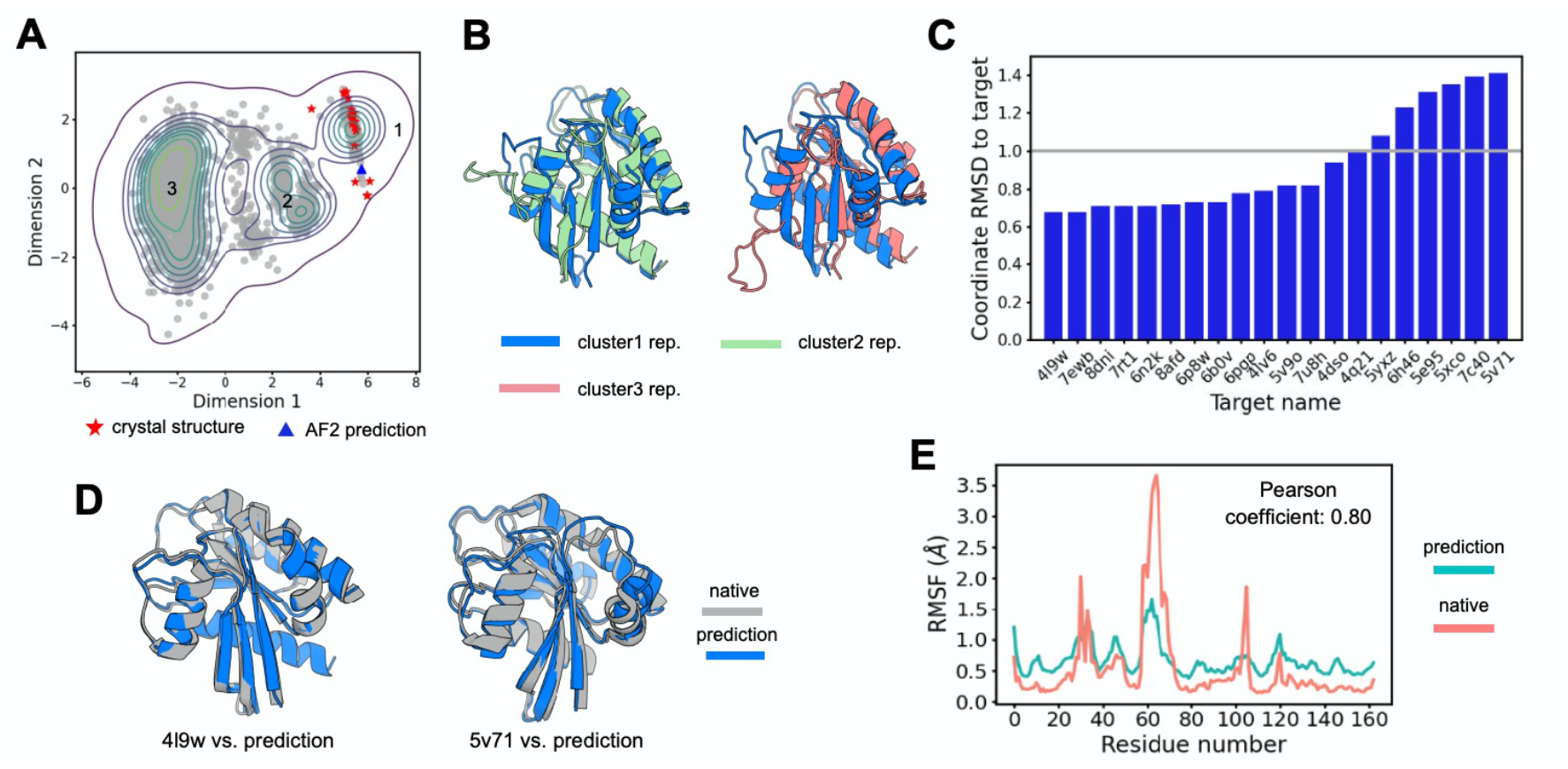
The conformational distribution of the K-Ras protein predicted by PVQD. A. The 2000 conformations predicted PVQD projected onto a plane obtained with Multi-Dimensional Scaling. The projections of 20 crystal structures and 2 AF2-predicted structures were also shown. The clusters formed by the predicted structures are numbered from 1 to 3. B. The superimpositions of the representative structure of cluster 1 (colored in blue) with the representative structures of clusters 2 (colored in green) and 3 (colored in salmon). C. The RMSDs of the experimental structures from the respectively most similar predicted structures (the smallest RMSDs). D. The superimpositions of the experimental and predicted structure pair with the lowest smallest RMSD (4l9w) and of the pair with the highest smallest RMSD (5v71). E. The residue-wise Root-Mean-Square Fluctuations (RMSFs) of the predicted structures in cluster 1 (colored in green) and the same RMSFs between the experimental structures (colored in salmon). The Pearson correlation coefficient is indicated.

Fig. 6A shows that the PVQD predicted structures for the D-allose binding protein ^29^ form three clusters. The representative structures of each cluster are shown in Fig. 8B. The two experimental structures and the two AF2-predicted structures all fall into cluster 1. The two experimental structures display inter domain motions with a TM-score of 0.72 (Fig. 8C), while their TM-scores to the respectively most similar predicted structures (the highest TM-scores) were 0.96 and 0.97, respectively (Fig. 8C). Fig. 8D shows that the structure variations between the predicted structures correspond to the same large inter-domain motions as the experimental structures. For the structural variations within the domains, the residue-wise structural fluctuations displayed by the predicted structures and the residue-wise deviations between the experimental structures are highly correlationed (Figures 8E and 8F).

**Fig 8.**
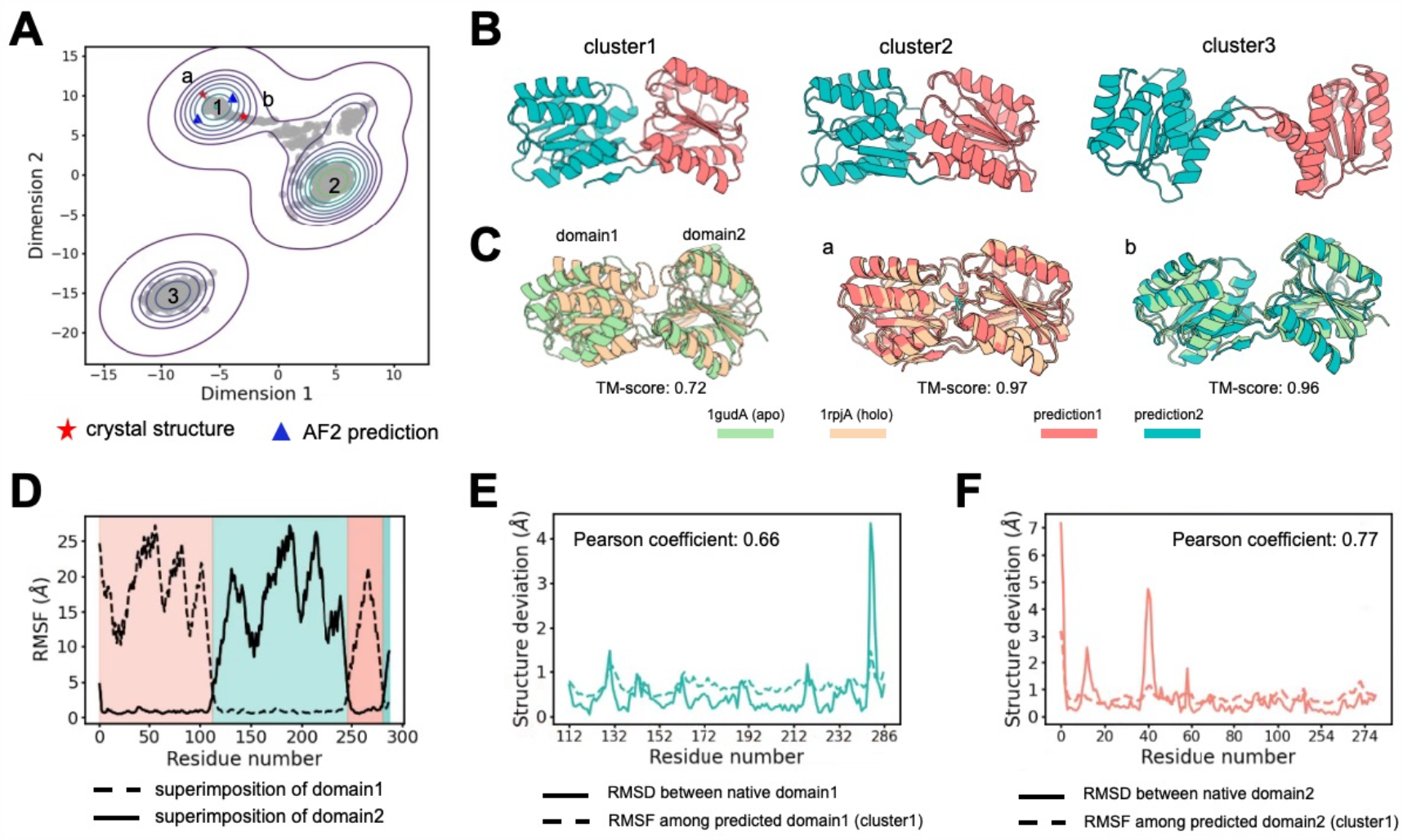
The conformational distribution of the D-allose binding protein predicted by PVQD. A. The 2000 conformations predicted PVQD projected onto a plane obtained with Multi-Dimensional Scaling. The projections of 2 experimental structures (a: 1gudA; b: 1rpjA) and 2 AF2-predicted structures were also shown. The clusters formed by the predicted conformations were numbered from 1 to 3. B. The representative structure of each cluster is displayed. The D-allose binding protein comprises two structural domains: domain 1 (residues 0-112 and 246-280, colored in green); and domain 2 (residues 112-246 and 280-287, colored in salmon). C. Left: the superimposition between two experimental structures. Middle: the superimposition between the experimental structure 1rpjA and its most similar (of the lowest RMSD) predicted structure. Right: the superimposition between the experimental structure 1gudA and its most similar (of the lowest RMSD) predicted structure. The TM-scores between the superimposed structures are indicated. D. The residue-wise RMSFs of backbone atom positions. The solid (dashed) line was computed by superimposing the residues in structural domain 1 (2) of the predicted structures. The residue ranges spanned by domain 1 (2) were shadowed in salmon (green). E. The residue-wise RMSFs of backbone atom positions for domain 1 in the predicted structures in cluster 1 (dashed line) in comparison and the displacements of backbone atom positions between the experimental structures of domain 1 (solid line). The Pearson correlation coefficient is indicated. F. The same as E, but for the RMSFs and displacements of backbone atom positions for domain 2.

Fig. 9A shows that the PVQD predicted structures for the F1-ATPase ^30, 31^ form two clusters. Projected onto the MDS plane, the two experimental structures can be separately assigned to the two clusters while the structures predicted with AF2 are located out of both clusters. The two experiment structures displayed complicated non-local structural changes with a TM-score of 0.82, which their highest TM-scores to predicted structures are both 0.98 (Fig. 9B). Fig. 9C shows that the residue-wise deviations between the predicted structures representing the two cluster centers and those between the two experimental structures are highly correlated.

**Fig 9.**
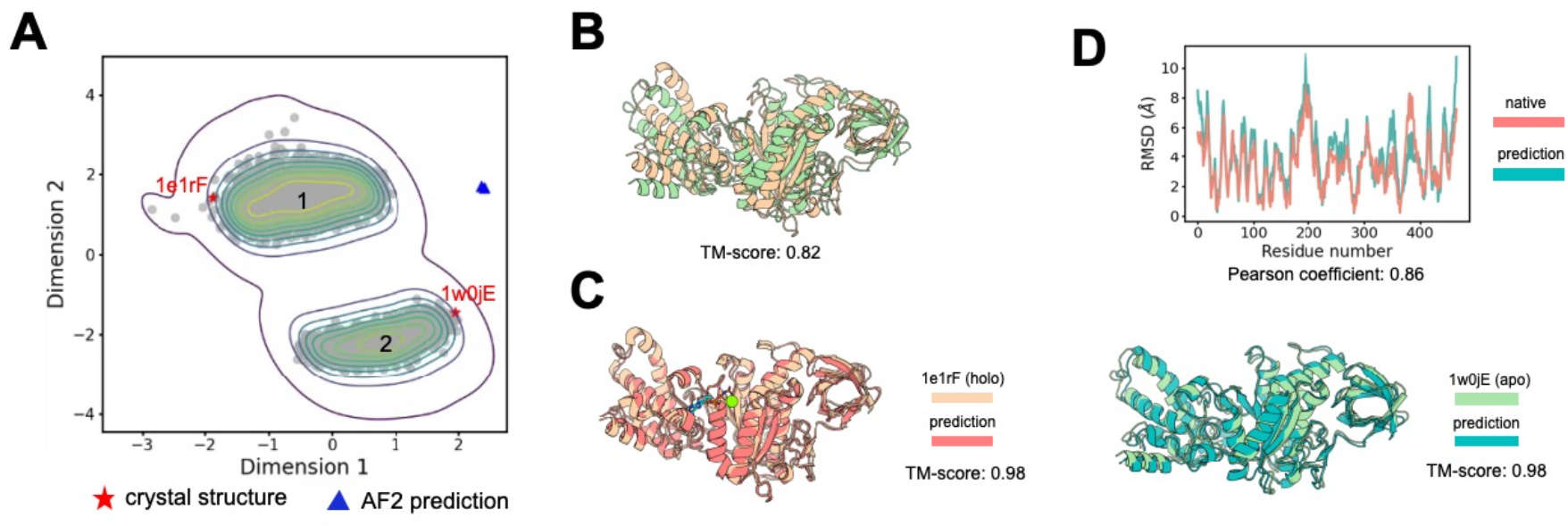
The conformational distribution of the F1-ATPase predicted by PVQD. A. The 2000 conformations predicted by PVQD projected onto a plane obtained with Multi-Dimensional Scaling. The projections of 2 experimental structures and 2 AF2-predicted structures were also shown. The clusters formed by the predicted conformations were numbered from 1 to 3. B. The superimposition between the two experimental structures. The TM-score between the superimposed structures is indicated. D The superimpositions between the experimental structures and its most similar (of the lowest RMSD) predicted structure. Left: for the experimental structure 1elrF (the holo state); right: for the experimental structure 1w0jE (the apo state). The TM-scores between the superimposed structures are indicated. C. The residue-wise displacements between the backbone atom positions of the representative predicted structure of cluster 1 and the representative predicted structure of cluster 2 (green) compared with the same displacements between the two experimental structures (salmon).

Thus, for all the three examples’ proteins considered here, the experimentally observed alternative conformations are faithfully covered by the respective set of conformations generated by PVQD. The predicted structures identified the same regions of high structural variability as the experimental structures. These results support the use of PVQD for identifying dynamic regions of proteins and for predicting possible alternative structures due to conformational dynamics.

## Discussion

The latent space vector quantization approach used by the auto-encoder of PVQD were originally developed for processing images and audios ^32, 33, 16, 34, 35, 36, 37^. The purpose of this approach is to compress continuous, high-dimensional data with high-frequency, imperceptible details into a finite set of discrete tokens (here the structural codes) embedded in a low-dimensional latent space. This results in significant dimension reduction and smoothening of the data distribution, facilitating the efficient learning of different generative models for various downstream applications. By performing sampling in a low-dimensional manifold without the need for treating details in the original space, these models can generate data of high diversity ^38, 16, 36, 37^. In the complete end-to-end model, these details are recovered by a well-trained decoder.

We note that in PVQD, the decoder recovers structural details by treating the structure codes at different positions not independently but cooperatively. For example, one structure code alone does not even uniquely determine the secondary structure state (Fig. 10). This indicates that the decoder plays substantial roles in the end-to-end generation of complicated structures, lessening the burdens on the diffusion-based generator.

**Fig 10.**
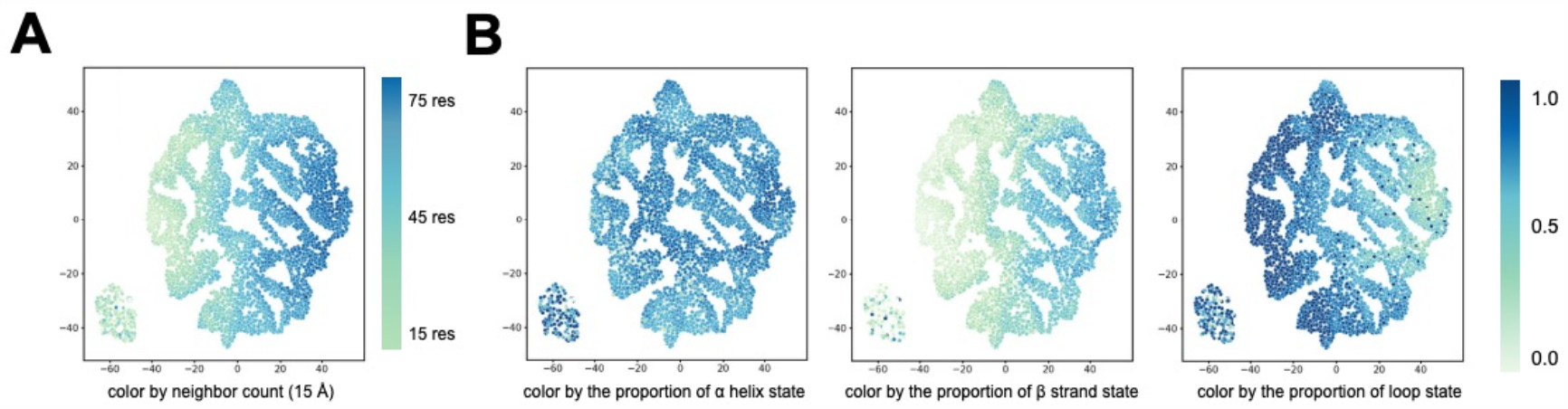
Visualization of the quantization vectors in the codebook of the auto-encoder. A. Two-dimensional projections (obtained with t-Distributed Stochastic Neighbor Embedding or t-SNE) of the quantization vectors in the codebook. Each projected point is colored according to the mean value of the count of neighboring residues (15 Å radius) averaged over the set of non-redundant natural protein structures (sequence identity cutoff 40% and X-ray structure resolution cutoff 2.0 Å). B. The same as A, but with the points colored according to the proportions of α helix state (left), β strand state (middle) and loop state (right).

Compared with protein structure design models using diffusion directly in the structural space ^8, 9, 10, 6^, PVQD shows an enhanced ability to generate structures of higher compositions of β-strands and longer loops. These structures, while being still designable according to computational metrics, may be intrinsically less rigid and can host richer conformational dynamics than structures designed with previous methods. As conformational dynamics is essential for important protein functions such as enzyme catalysis and allosteric regulations, it will be of high interest to verify that such PVQD designed structures and their conformational dynamics can indeed be realized in wet experiments.

Predicting conformational distributions from amino acid sequence is also of high interest for investigating the functional dynamics of natural proteins. Although there were a few recent studies examining the use of existing structure prediction networks (especially AlpahFold2) for this purpose by, *e.g*., perturbing the input multiple sequence alignments ^14^, sampling-based models such as DDPMs offer a more natural approach. We have demonstrated that besides achieving comparable performances for single amino acid sequence-based structure prediction as several current state-of-the-art methods, PVQD could be used to sample alternative structures due to protein conformational dynamics. For a majority of proteins in the benchmark set contained here, PVQD produced different predicted structures that approximated different experimental structures with errors smaller than the differences between the experimental structures themselves. Moreover, for three example proteins exhibiting different types of dynamic changes, the PVQD predicted structures and the experimental structures indicated almost the same high variability regions. As PVQD achieved this without using protein-specific models or training with data like molecular dynamics trajectories, we expect it to be widely applicable for exploring the conformational dynamics of natural proteins to understand their function and regulation.

## Methods

### The auto-encoder of PVQD

The encoder module of the auto-encoder of PVQC uses a graph-based Geometry Vector Perceptron (GVP) ^39^ to encode and transform the structural context of a central residues surrounded by its 30 nearest neighbor residues. Each node of the graph corresponds to a residue. The central residue node is connected to each nearest-neighbor residue node by an edge. The node features of residue *i* include the unit vectors along the directions of *Cα*_*i+1*_*– Cα*_*i*_, *Cα*_*i-1*_*– Cα*_*i*_ and 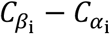. The edge features between two spatially neighboring residues i and j include their relative sequence position encoded with a sinusoidal scheme ^40^, the distance ‖*Cα*_*j*_ *– Cα*_*i*_*‖*_*2*_encoded with a set of Gaussian radial basis functions, and the unit vector along the direction of *Cα*_*j*_ *– Cα*_*i*_. The three-dimensional vectors are represented in the local coordinate frame defined based on the backbone N, *Cα*, and C atoms of the central residue and are thus SE(3)-invariant. The graph is transformed by a GVP network using four repeated Transformer encoder blocks to update the concatenated node features. From the last layer of the GVP, the output concatenated vector associated with the central residue node is extracted and projected into a 16-dimensional vector, which is used as a latent space representation of the residue-wise structural context.

The codebook is formed through vector quantization of the above latent space into discrete codes. Given the codebook size (*i.e*., the number of codes) K as a hyperparameter, a set of representative vectors *e*_*k*_, *k ∈ 1 … K* in the latent vector space are learnt. Any vector *v* in the latent space is mapped to (or approximated by) the closest *e*_*k*_, namely,

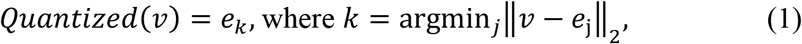

The codebook together with the encoder maps the backbone structure of a protein (denoted as *x*) to a code sequence *q = {q*_*1*,_*q*_*2*,_*…,q*_*N*_*}*, in which each *N* is the sequence length and for every *i ∈ 1 … N, q*_*i*_ *∈ 1 … K* stands for the integer code index at position *i*. Denoting the corresponding set of latent space vectors as 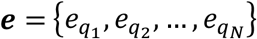, we denote the transformation from *x* to ***e*** as ***e****=E(x)*.

The decoder of PVQD reconstructs *x* from ***e*** with repetitive Transformer-based blocks and Invariant Point Attention (IPA) blocks ^10, 2^. The sequence of quantization vectors was initially processed with four repeated Transformer blocks. From the resulting residue-wise representation, a residue pairwise representation was derived by first splitting the channel dimensions of each residue into two groups, namely,

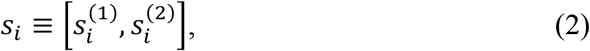

where *s*_*i*_ is the representation of residue *i*, and then recombining the channels for each pair of residues, *i.e*.,

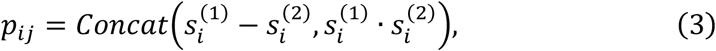

where *p*_*ij*_ is the representation of the residue pair *i* and *j*. The residue-wise representation and the pair-wise representation were further processed and updated with repeated IPA-Transformer blocks to generate the three-dimensional backbone structure. We denote the entire transformation from ***e*** as 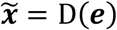, in which 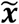 stands for the reconstructed structure.

The encoder network *E*(*·*), the codebook ***e*** and the decoder network *D(·*) are learnt by optimizing the reconstruction losses and vector quantization losses on training backbones. For a backbone *x*, the total loss is

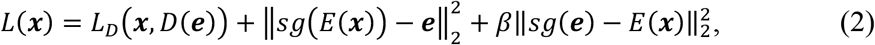

in which the first term *L*_*D*_(*x,D(e*)) stands for the structure reconstruction loss, and the second and third terms represent the vector quantization losses. With the operator *sg* referring to the stop-gradient operation that blocks the back propagation of error gradients, the second term leads to the optimization of the codebook ***e***, while the third term (the commitment term) leads to the optimization of the encoder. The hyperparameter *β*(chosen to be 0.25 in this work) controls the relative weights.

The total structure reconstruction loss is composed of the Frame Align Point Error (FAPE) loss as used by AF2 ^2^ supplemented with four auxiliary losses: a distogram classification loss (*CE*_*dist*_), a backbone conformation loss (*MSE*_*Violation*_), a covalent geometry violation loss (*MSE*_*BBconf*_) and an amino acid type recovery loss (*CE*_*aatype*_). That is,

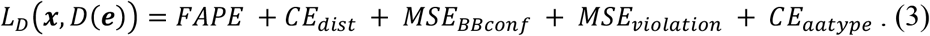

Here, the amino acid types were predicted from the residue-wise representation in the last IPA block of the structure decoder, while the categorized inter-atomic distances were predicted from the pair representation of the same IPA block.

We note that while the squared error losses FAPE, *MSE*_*BBconf*_ and *MSE*_*Violation*_ are based on the reconstructed three-dimensional structures, the cross-entropy losses *CE*_*dist*_ and *CE*_*aatype*_ are computed with separate classifiers decoding respectively inter-atomic distance categories and amino acid types based on the last internal layer of the structure decoder.

We trained the auto-encoder using the structures of natural proteins from the PDB database until Apr 30, 2020. The available set of non-redundant natural structures (selected with the PISCES server ^21^ with a sequence identity cutoff of 70% and X-ray structure resolution cutoff 3.5 Å) were randomly split into a training set (99%) and a validation set (1%). During the training process, we selected the natural structures with less than 640 residues to train the auto-encoder. More details about training of the auto-encoder are shown in Table 2. The validation set was used during training for comparing hyper-parameters and monitoring the convergence of losses.

**Table 2.**
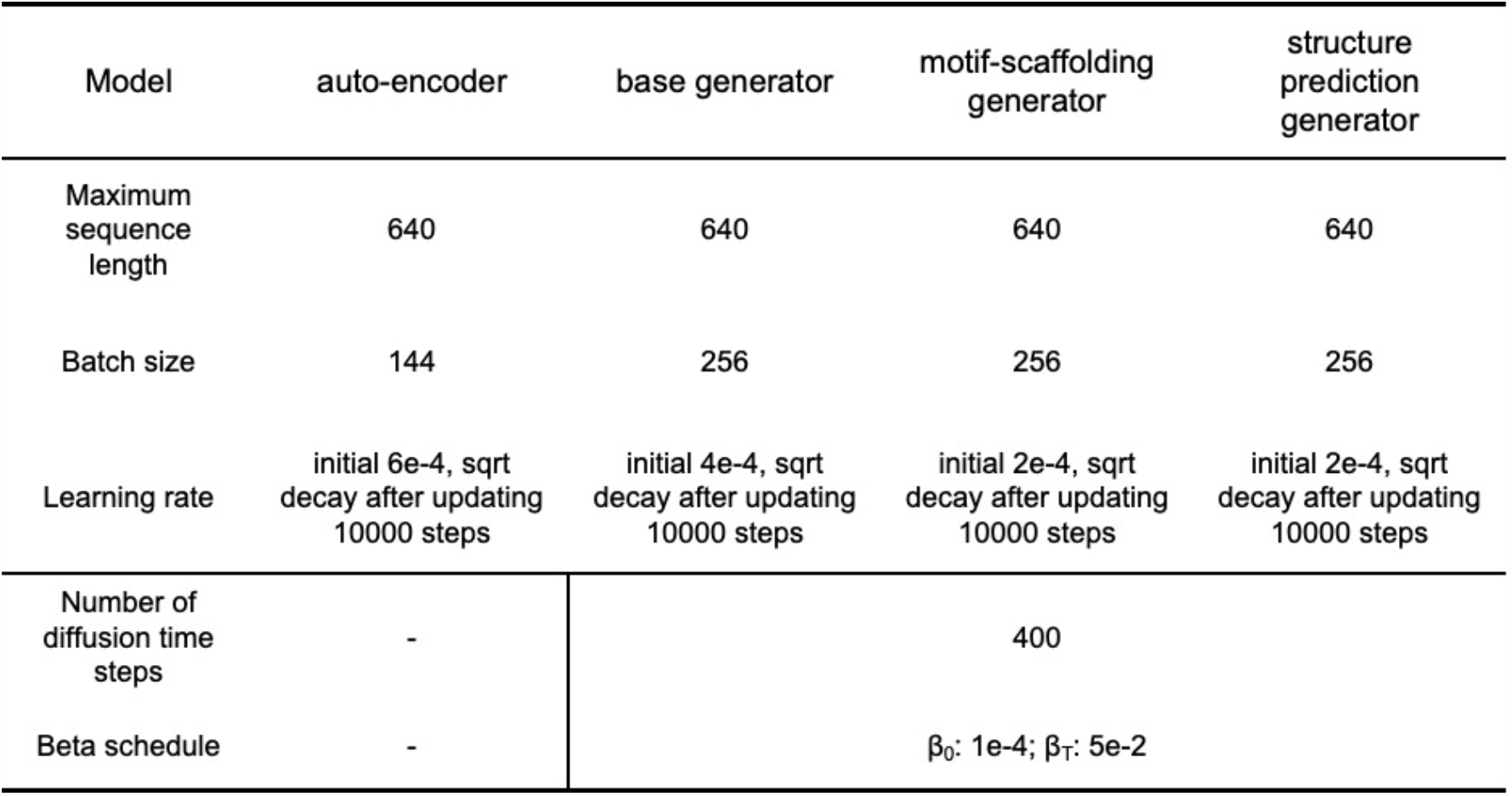
Parameters for training PVQD.

### The diffusion-based generator of PVQD

We modeled the joint distribution of the latent space vectors encoding backbone structures with a denoising diffusion probabilistic model (DDPM). In DDPMs, a forward Markovian diffusion process of *T* time steps are used to gradually introducing Gaussian noise into the true data, while a network is trained to perform the inverse denoising process to recover the true data. Here we used the denoising network architecture of Diffusion Transformers ^20^. The module was composed of 24 repeated Transformer blocks. The time step embedding is incorporated through the adaptive Layer Norm (AdaLN) modules. Through denoising diffusion from Gaussian random noise, a sequence of the latent space vectors is generated by the diffusion module, which is subsequently mapped to a sequence of the quantized vectors, and then decoded into a three-dimensional backbone structure as in the auto-encoder.

The generator was trained by minimizing the Mean Square Error (MSE) between the generated latent space vectors and the quantized vectors of the native backbone. The MSE loss computed on the validation set was monitored during training, and convergence was reached after approximately 200k training updates.

The DDPM trained for unconditional generation was used as a base version, which was fine tuned to develop the PVQD model for the conditional generation of backbones containing pre-defined partial backbone structures (*i.e*., motif-scaffolding). In this model, a conditioner using the same GVP architecture as the encoder module of the auto-encoder is used to encode the pre-defined partial structure (the parameters of the conditioner were also initialized with their corresponding values in that module during training). The resulting embedding is added with the time step embedding and incorporated into diffusion through the AdaLN modules as in the Diffusion Transformers ^20^. Additionally, after the self-attention module in each Transformer block, a Multi-Head Cross-Attention between the embedding of the structural conditions and the diffusing latent space vectors was performed to align the two embedded representations. To fine tune the model, the training natural backbones with the structures of 40% to 90% residues masked were used as conditions of predefined partial structures. The masked residues were either multiple segments of 3 to 7 residues, or a single long segment, or a set of residues that are spatial neighbors surrounding a randomly selected residue. The same convergence criterion as for training of the base model was used for the fine-tuning process.

The PVQD model for structure prediction from single amino acid sequence was also developed by fine tuning the base DDPM for generating backbones with the condition of given amino acid sequences. The amino acid sequence was embedded with the pre-trained Evolutionary Scale Modelling or ESM network (650M parameters) ^23^ and integrated (with frozen parameters) into the Diffusion Transformers as described above. The final model was trained on the same set of training proteins as described above. For evaluation, a model for predicting the conformational distributions of the 91 benchmark proteins were specially trained by removing all proteins with more than 70% of sequence identity with any benchmark protein from the training set.

## Acknowledgments

We thank the support by the Huawei MindSpore team.

